# Control of Intracellular Molecular Networks Using Algebraic Methods

**DOI:** 10.1101/682989

**Authors:** Luis Sordo Vieira, Reinhard C. Laubenbacher, David Murrugarra

## Abstract

Many problems in biology and medicine have a control component. Often, the goal might be to modify intracellular networks, such as gene regulatory networks or signaling networks, in order for cells to achieve a certain phenotype, such as happens in cancer. If the network is represented by a mathematical model for which mathematical control approaches are available, such as systems of ordinary differential equations, then this problem might be solved systematically. Such approaches are available for some other model types, such as Boolean networks, where structure-based approaches have been developed, as well as stable motif techniques.

However, increasingly many published discrete models are mixed-state or multistate, that is, some or all variables have more than two states, and thus the development of control strategies for multistate networks is needed. This paper presents a control approach broadly applicable to general multistate models based on encoding them as polynomial dynamical systems over a finite algebraic state set, and using computational algebra for finding appropriate intervention strategies. To demonstrate the feasibility and applicability of this method, we apply it to a recently developed multistate intracellular model of E2F-mediated bladder cancerous growth, and to a model linking intracellular iron metabolism and oncogenic pathways. The control strategies identified for these published models are novel in some cases and represent new hypotheses, or are supported by the literature in others as potential drug targets.

Our Macaulay2 scripts to find control strategies are publicly available through GitHub at https://github.com/luissv7/multistatepdscontrol.

## 1 Introduction

Modification or differential regulation of intracellular networks, either at the gene, protein, or metabolite level, can result in an altered cellular phenotype. Being able to perform targeted modifications for this purpose may be desirable for several reasons, whether to alter bacterial metabolism for industrial production [38] or to mitigate properties of a tumor cell [16,17,59]. Systematic approaches to the identification of such targeted modifications are therefore of considerable importance. Generally, this is accomplished through the use of mathematical models as discovery tools. In addition to systems of differential equations, an increasingly common modeling framework are time- and state-discrete models, such as Boolean networks and their various generalizations. These provide only semi-quantitative information but are more easily constructed, since they do not require quantitative kinetic information, and they can sometimes be more intuitive for the experimentalist. A drawback of discrete models is that their underlying mathematical theory, in particular control theory, is not yet well-developed, and the present paper makes a contribution to this body of work.

Given a mathematical model of an intracellular regulatory network, one commonly associates the possible phenotypes of the cell with the attractors of the model, an idea that can be traced back to Waddington [50,42] and Kauffman [23,21]. For example, the steady states in [37], discussed in more detail below, correspond to proliferative, apoptotic, or growth-arrest phenotypes of a cancer cell. In [3], the steady states of the model correspond to the observed altered iron metabolism phenotypes in a breast epithelial cell with and without a certain RAS mutation. We present here a method to systematically identify modifications to a model that can change the attractor landscape in prescribed ways. Such modifications consist of, e.g., deactivating a node or modifying the effect of an edge in the model’s wiring diagram, a graph-theoretic representation of the functional dependencies of the different model variables. In the case of gene regulatory or signaling networks, this could be accomplished, for instance, through a compound that blocks the protein corresponding to a particular gene.

We focus on the mathematical modeling framework of multistate discrete networks. These can be defined as dynamical systems that are discrete in time as well as in variable states. More formally, consider a collection *x*_1_,…, *x_n_* of variables, each of which can take on values in a finite set *X*_1_,…, *X_n_*. Let *X* = *X*_1_ × ⋯ × *X_n_* be the Cartesian product. A discrete dynamical system in the variables *x*_1_,…, *x_n_* is a function

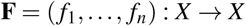

where each coordinate function *f_i_* : *X* → *X*_1_ represents how the future value of *x*_1_ depends on the present values of all the variables. If *X*_1_ = {0, 1} for all *i*, then each *f_i_* is a Boolean rule and **F** is a Boolean network.

Discrete networks defined in this way can be represented in the richer mathematical framework of *polynomial dynamical systems* (PDSs) [48,19], as can models in other common frameworks, such as Boolean networks [1], logical regulatory graphs [2], or multistate networks [37,3,11,45]. For instance, the mathematical tools associated with PDSs allow for the computation of all steady states and cycles up to a certain length of a system as the solutions to a system of polynomial equations (in a suitably chosen finite field) without explicit simulation of the entire state space. Recently, we have used these mathematical tools to construct a method that rigorously computes modifications for the control of Boolean networks that can help avoid regions of the state space or create new steady states or cycles [32]. The method was applied to a mathematical model of cellular response to DNA damage [4] and a model of large granular lymphocyte apoptosis escape [60]. However, an increasing number of discrete mathematical models in this context are not Boolean, e.g., [37,3,11,45], so that a more general method is desirable, and such a generalization is the focus of this paper.

We note that there are several published control methods that do not rely on the PDS representation of discrete models, such as Stable Motifs [55], Feedback Vertex Sets [57], Minimal Hitting Sets [49,25], and several others [36,26,35,14,56]. Our method provides a flexible control framework that, for instance, allows for the identification of controllers for creating new (desired) steady states, a feature that other methods do not allow.

It also extends our method for Boolean networks [32,31] to multistate networks, and thus broadening the scope of use of the PDS representation of discrete models.

As our method uses polynomial algebra over a finite field, all network nodes need to take values in a common finite field, in particular, all nodes need to have the same number of possible values. In many published models, however, different nodes take on different numbers of states, and this number generally does not allow the imposition of a finite field structure (for which the number is required to be a power of a prime number); see, e.g., [37,56]. As part of the algorithm in this manuscript, we present a method to convert models with a general number of mixed discrete states into a model that satisfies the computational algebra requirements, without changing the model’s steady states, and which is not equivalent to the well-known reduction to a Boolean network that adds new nodes to the network, as done in [56].

We demonstrate the power and versatility of this method by applying it to two recently published multistate network models. One is a model of bladder cancer response to different stimuli, including DNA-damage, EGFR, FGFR3, and growth inhibitors [37]. The method can find combinatorial interventions that block proliferative steady states. The second is an intracellular iron network model in breast epithelial cells presented in [3]. We identify interventions to recover basal expression of the iron export protein of a malignant breast epithelial cell with RAS over-expression. These can be viewed as predictions to be validated experimentally.

## 2 Methods

As the first step of the method, a multistate discrete network is represented in an algebraic framework. In the process, we provide a general procedure to extend any multistate discrete network to a polynomial dynamical system. We refer the reader to the following books [7,27] for the basics of finite fields and the basics of computational algebra.

### 2.1 Discrete dynamical systems

In this manuscript, we consider discrete variables *x*_1_,…, *x_n_*, each taking on values in a finite set *X*_1_,…, *X_n_*.

For the purpose of exploiting the algebraic properties of discrete functions, it is assumed that the variables *x*_1_,…, *x_n_* take on values on a finite field 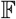. Then, using the fact that any discrete function 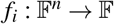 can be represented as a polynomial on {*x*_1_,…, *x_n_*}(see *e.g*. [22,27]), that is 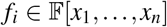, the discrete network can be represented as

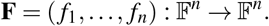

The discrete network can thus be represented as a *polynomial dynamical system* (PDS) [48,19]. We describe how to convert a mixed-state model into a PDS in the Appendix, Section 7.1. We also provide a concrete example of how to carry out this procedure when the update rules are written as Boolean expressions of conditions in the Appendix, Section 7.2.

Given a discrete network **F** = (*f*_1_,…, *f_n_*), we can define its wiring diagram 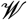 to be the directed graph with *n* nodes *x*_1_,…, *x_n_* associated to **F**, such that there is a directed edge in 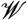 from *x_j_* to *x_i_* if *x_j_* appears in *f_i_*, and there exist

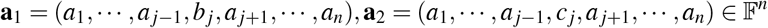

such that

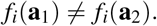

In other words, the value *f_i_* takes on depends on the values of *x_j_*.

The dynamics of discrete networks are given by the difference equation *x*(*t* + 1) = **F**(*x*(*t*)); that is, the dynamics are generated by iteration of **F**. More precisely, the dynamics of **F** are given by the state space graph S, defined as the graph with vertices in 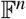 which has an edge from 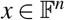 to 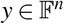 if and only if *y* = **F**(*x*). In this context, the problem of finding the states 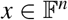 where the system will get stabilized is of particular importance. These special points of the state space are called attractors of a discrete network and these attractors may include steady states (fixed points), where **F**(*x*) = *x*, and cycles, where **F**^*r*^(*x*) = *x* for some integer *r* > 1. Attractors in network modeling might represent a differentiated cell type [24] or a cellular state such as apoptosis, proliferation, or cell senescence [20,41]. Identifying the attractors of a system is an important step towards system control. For example, a steady state might represent a cellular phenotype characterized by low expression of an iron exporter [3], a particularly deleterious phenotype in several cancers [46].

### 2.2 Control actions

In this manuscript, we focus on controlling the dynamics of a multistate network by avoiding undesirable steady states, creating new steady states, or avoiding regions in the state space, accomplished by modifications to the wiring diagram of **F**. This is an extension of our previous control methods applicable to Boolean networks [32].

We consider two types of control actions: 1. deletion (or constant expression) of edges and 2. deletion (or constant expression) of nodes. An edge deletion represents the experimental intervention that prevents a regulation from happening. These actions can be achieved by the use of therapeutic drugs that target a specific gene interaction [4]. Constant expressions could also help to drive the system into a more desirable state [39].

#### 2.2.1 Encoding control actions in multistate networks

In the Boolean setting, the deletion of an edge in the wiring diagram was implemented by setting an input to zero so that the interaction of that input (represented by an edge) was being silenced. For the multistate case, the silencing of the interaction will be applied whenever the control variable is within a range of values of the possible discrete values. For expository purposes, we use a simple function taking the value 1 on the singleton set {0} and 0 on the set 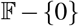.

**Definition 1 (Edge control for multistate networks)** Consider the edge *x_i_* → *x_j_* in the wiring diagram 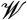 and let 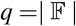.

For 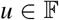, the control of the edge *x_i_* → *x_j_* consists of manipulating the input variable *x_i_* for *f_j_* in the following way:

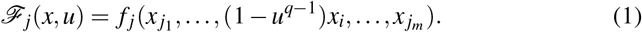

For each value of 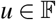 we have the following control settings:

– For *u* ≠ 0, 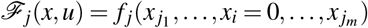. This is the case when the control is active and the action represents the removal of the edge *x_i_* → *x_j_*.
– For *u* = 0, 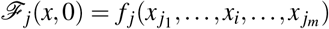. That is, the control is not active.

Similarly, for node deletion in the multistate setting we have the following definition.

**Definition 2 (Node control for multistate networks)** Consider the node *x_i_* in the wiring diagram 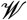 and let 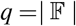. The function

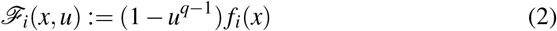

encodes the control of the node *x_i_* because for each possible value of 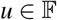 one has the following control settings:

– For *u* ≠ 0, 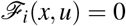. This action sets the function of *x_i_* to zero. For instance, this can represent the knock-out of gene *x_i_* or blocking the synthesis of a protein *x_i_*.
– For *u* = 0, 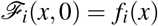. That is, the control is not active.

We note that in Equations 1-2 we only consider edge and node deletions. In general, one could consider setting edges and nodes to a constant value within 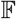.

#### 2.2.2 Generating new steady states

Suppose that 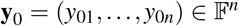 is a desirable cell state (for instance, it could represent the state of cell senescence in a cancer model) but is not a steady state, i.e., **F**(**y**_0_) = **y**_0_. The problem, then, is to choose a control 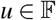 such that 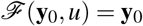. We now show how this can be achieved in our framework.

After encoding the multistate network with control as a PDS

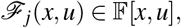

we consider the system of polynomial equations in the *u* parameters:

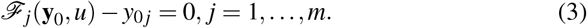

#### 2.2.3 Destroying existing steady states or, in general, blocking transitions

Suppose that 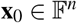 is an undesirable steady state of **F**(*x*), that is, **F**(**x**_0_) = **x**_0_ (for instance, it could represent a tumor proliferative cell state that needs to be avoided). The goal here is to find a set of controls such that 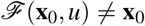. More generally, one may want to avoid a transition between two states **x**_0_ and **z**_0_. That is, we want to find controls such that 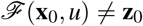. To solve this problem consider the following equation,

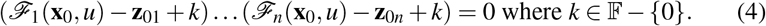

Equation 4 defines a system of polynomial equations in the *u* parameters. It can be shown that 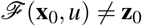 if and only if Equation 4 is satisfied.

#### 2.2.4 Blocking regions in the state space

We now consider the case where we want the dynamics to avoid certain regions of the state space. For example, if a particular value of a variable, 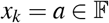, activates an undesirable pathway, or is the signature of an abnormal cell state, then we want all steady states of the system to satisfy *x_k_* ≠ *a*. In this case, we consider the system of equations

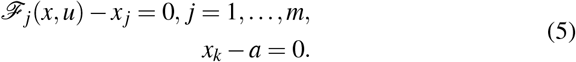

Note that in contrast to previous sections, we are now using variables for *x* instead of specific values. Since the steady states with *x_k_* = *a* are to be avoided, we want to find controls *u* for which Equation 5 has no solution.

#### 2.2.5 Identifying control targets

In each case of the tasks above we obtain a system of equations (or a single equation) that we need to solve to find the appropriate controls. This can be done using computational algebra tools. For instance, we can compute the Gröbner basis of the ideal associated with Equation 3, see [6],

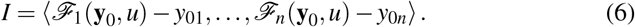

The computation of a Gröbner basis allows us to read off all controls as the solutions to the system of equations. Furthermore, the algebraic approach can detect combinatorial control actions such as control by the synergistic combination of more than one action; see Section 3 for examples.

## 3 Results

We first apply the control methods to the mathematical intracellular bladder cancer model in [37]. Then we will apply our methods to the intracellular iron network model in breast epithelial cells in [3]. As a sample control problem, we first show how we can find control strategies for blocking proliferative steady states by blocking interactions and nodes. Then we will identify interventions to recover a desirable fixed point in a malignant breast epithelial cell with RAS over-expression.

### 3.1 Description of E2F-mediated growth model

In [37], the authors present a generic network centered around how a cell in response to stimuli such as the growth factors FGF3, EGF3, and nodes representing growth inhibition and DNA damage ends up in different states such as a proliferative state, apoptotic state, or growth arrest. The model includes several of the cyclins which regulate cell cycle progression by interactions with cyclin-dependent kinases, as well as the E2F family of transcription factors, which are released from pRB inhibition in the G1/S cell cycle state and control the transcription of several factors relevant to complete the cell cycle. The model is a multistate model with a total of 30 nodes, where 25 nodes are binary and 5 nodes are ternary. Four nodes serve as input, namely FGF3, EGF3, Growth Inhibition and DNA damage. Three nodes serve as output, namely Proliferation, Apoptosis, and Growth Arrest; see Figure 1 for the wiring diagram.

**Fig. 1:**
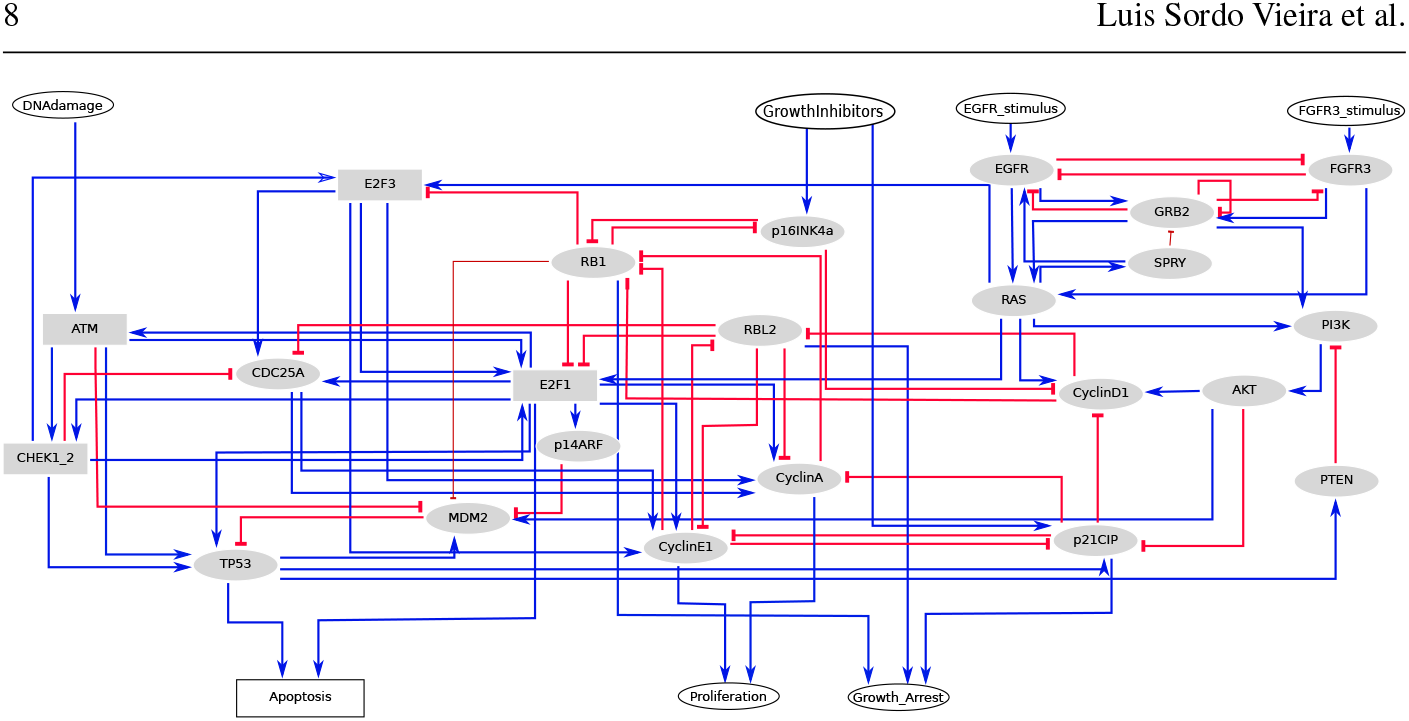
Wiring diagram for the model of bladder tumorigenesis published in [37]. This figure was generated using GINsim [33]. The red arrows represent inhibitory interactions and the blue arrows represent interactions with an activation/positive effect.

This mathematical model can then be used to see which inputs lead to a cancerous (proliferative) phenotype, or to generate hypotheses on knockdowns, and/or overexpression of proteins that will evade the proliferative steady state. Due to the size and connectivity of the model, it quickly becomes apparent that predicting emergent behaviors from the blocking of interactions and nodes in the model is not easily done.

#### 3.1.1 Converting the model into a polynomial dynamical system over 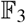

In order to use the PDS framework, we first convert the model in [37] into a polynomial dynamical system over 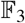. To do so, we expand the state space to 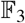 but in such a way that the steady states do not change. See the appendix for details. The correspondence between variables in the polynomial dynamical system and nodes in Figure 1 is given in Table 1. We used a Docker image of Macaulay2 v: 1.14 [15] and a script available in the Github repository for the conversion.

**Table 1:**
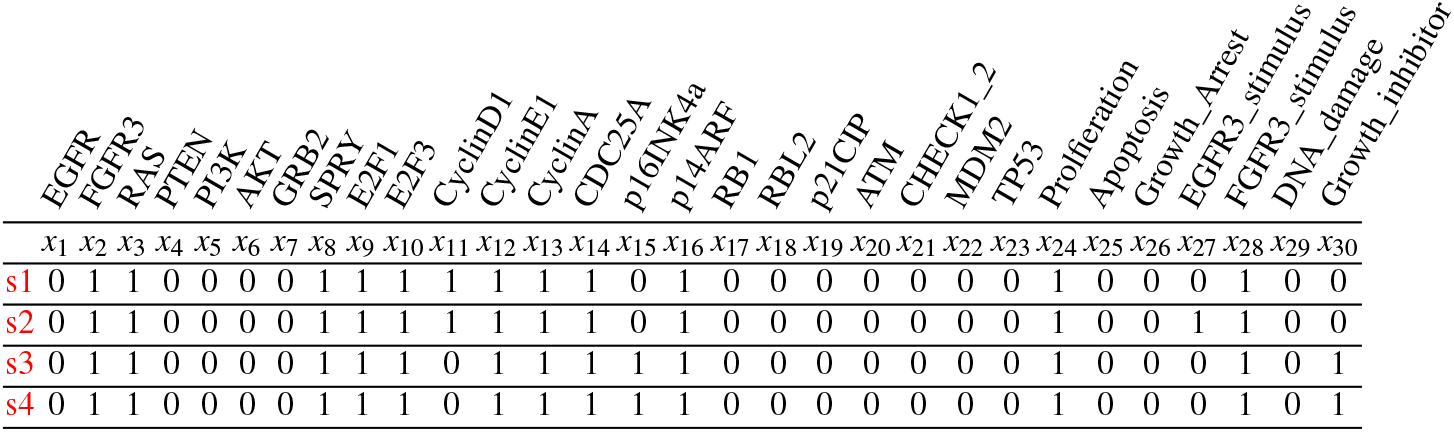
Steady states of the original model presented in [37] where Proliferation=1.

### 3.2 Avoiding proliferative steady states

In the original model (see Fig. 1), there are four steady states with proliferation equal to 1 (*x*_24_ = 1, Table 1), that is, steady states in which the cells proliferate. It is interesting to note that these steady states have TP53, PTEN, and p21CIP activity at 0, which are all well-known tumor supressors. As mentioned before, steady states correspond to cellular phenotypes. A proliferative steady state thus potentially corresponds to a cell undergoing uncontrolled proliferation. A natural biological question thus arises. How do we avoid such steady states?

Our control method predicts several possible control strategies, of which six result in non-proliferative steady states (Table 2). Additional control methods that destroy the original steady states can be combined to block all possible proliferative steady states.

**Table 2:**
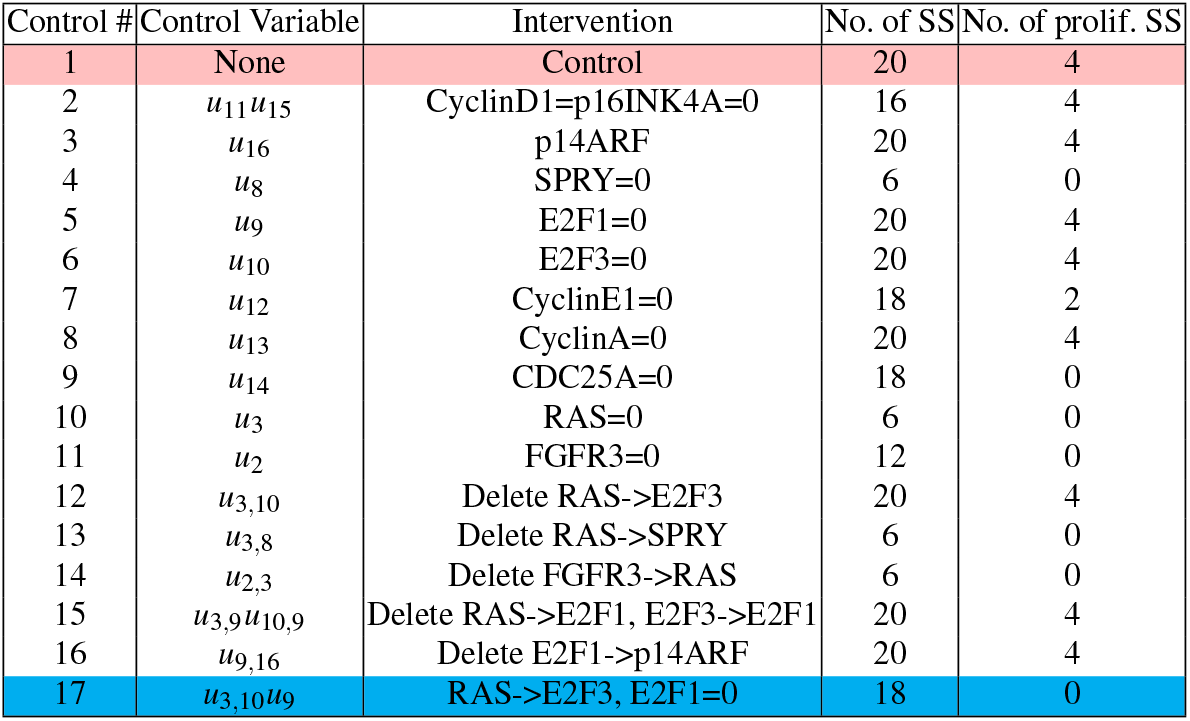
Controls for the model presented in [37]. The first row (pink) corresponds to the original model, where no control was applied. The last row (cyan) corresponds to combining Control #5 with Control #12.

| Control # | Control Variable | Intervention | No. of SS | No. of prolifer. SS |
| --- | --- | --- | --- | --- |
| 1 | None | Control | 20 | 4 |
| 2 | $u_{11}u_{15}$ | CyclinD1=p16INK4A=0 | 16 | 4 |
| 3 | $u_{16}$ | p14ARF | 20 | 4 |
| 4 | $u_8$ | SPRY=0 | 6 | 0 |
| 5 | $u_9$ | E2F1=0 | 20 | 4 |
| 6 | $u_{10}$ | E2F3=0 | 20 | 4 |
| 7 | $u_{12}$ | CyclinE1=0 | 18 | 2 |
| 8 | $u_{13}$ | CyclinA=0 | 20 | 4 |
| 9 | $u_{14}$ | CDC25A=0 | 18 | 0 |
| 10 | $u_3$ | RAS=0 | 6 | 0 |
| 11 | $u_2$ | FGFR3=0 | 12 | 0 |
| 12 | $u_{3,10}$ | Delete RAS->E2F3 | 20 | 4 |
| 13 | $u_{3,8}$ | Delete RAS->SPRY | 6 | 0 |
| 14 | $u_{2,3}$ | Delete FGFR3->RAS | 6 | 0 |
| 15 | $u_{3,9}u_{10,9}$ | Delete RAS->E2F1, E2F3->E2F1 | 20 | 4 |
| 16 | $u_{9,16}$ | Delete E2F1->p14ARF | 20 | 4 |
| 17 | $u_{3,10}u_9$ | RAS->E2F3, E2F1=0 | 18 | 0 |

It is often difficult to generate drugs that target specific proteins, as is often the case of ‘undruggable’ targets [8]. Moreover, the knockdown of a node could be attained by knocking out a particular gene, but proteins have multiple indispensable physiological actions, and thus a gene knockout might be lethal. It is thus of interest to also be able to affect interactions between two products, which in our context, can be done by targeting edges in the wiring diagram.

To show how this can be done, consider the set of variables

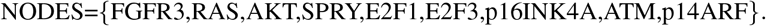

We consider the blocking of edges if and only if source and target nodes are both in the set NODES. We also allow node control for nodes *x*_1_ to *x*_23_.

We encode the control of edges and nodes as in Section 2.2.1, and find all solutions to the system encoded by the ideal 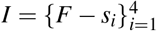. It is unfeasible to solve these equations by hand, so we use computer algebra. We compute generators for the Gröbner basis of *I*, and look at the generators comprised of *us* alone. The computation of the generators of the Gröbner basis took an average of 3.3653s to compute, with a standard deviation of 0.07636615. We get the following control variables

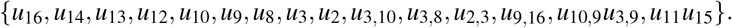

In order to avoid the steady states, we can choose any of these generators to not have value 0. The *u_i_*s correspond to nodes that are candidates for a knock-down, and the *u_i, j_*s correspond to interactions from node *x_i_* to *x_j_* to block. The product of two control variables indicates that both controls should be applied simultaneously. For example, *u*_10,9_*u*_3,9_ indicates that the interactions from E2F3 and RAS to E2F1 should be simultaneously inhibited. We present all generated controls in Table 2 with the number of resulting steady states, as well as with the number of proliferative steady states.

It is worth exploring mixed edge-node controls to attempt to destroy all proliferative steady states. For example, Control # 17 in Table 2 is derived by mixing Control # 12 with Control # 5.

Importantly, we see that instead of targeting RAS (Control #3), which has proven to be a formidable challenge and considered a holy grail of cancer therapeutics [43, 44], deleting the interaction between RAS and SPRY (Control #13) or FGF3 and RAS (Control #14) has the same effect (in terms of number of proliferative steady states and total steady states) as targeting RAS. Importantly, as RAS is the only node activating SPRY in this model, inhibiting the interaction between RAS and SPRY (Control #13) is equivalent to silencing SPRY, which has been shown to be beneficial in a xenograft model of rhabdomyosarcoma tumors [40]. Furthermore, it has been suggested that the FGF3 mediated RAS activation (Control #14) can lead to Vemurafenib resistance in melanoma cells [51]. Interestingly, inhibiting the CDC25 (Control # 9) family has been suggested as a potential therapeutic for triple negative breast cancer [28], and we see that knocking down CDC25A results in no proliferative steady states.

We thus see that we find some control strategies that destroy all proliferative steady states, some of which have shown promising results in different types of cancers. We also see that some control strategies are not completely effective (for example, knocking down E2F1 alone (Control #5) resulted in four new proliferative steady states, Table 2). However, the control candidates presented here quickly narrow down our list of possible targets. Furthermore, by using function composition, we can also set up control strategies to avoid cycles of a given length.

### 3.3 Algebraic methods applied to the Booleanized network model

The original model presented in [37] is naturally presented in a Boolean network model. It is natural to wonder how the algebraic methods of control compare if we apply them to the representation over 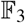 or to the original representation over 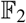. We applied the same methods of control to the network as explained in Section 3.2 (code to do this is in the Github repository). The computation of controls applied 10 times in a Dockerized image of Macaulay2 to the Booleanized model took an average of 6.3528 s with a standard deviation of 0.1718745 (including the running and removal of the Docker container), which is almost twice the time we had for the multistate case (AV=3.37 and SD=0.08). Notice that for the Booleanized network we add an extra node for each multistate node to represent the different possible levels. For example, E2F1 now becomes E2F1:1 and E2F2:2 to represent the possible values. In general, the number of variables to be added could rapidly grow and add computational complexity to the Groebner basis computation. Moreover, without Booleanization, the control nodes and edges are directly related to the wiring diagram of the biological system, making the results easier to interpret and more readily actionable.

### 3.4 Comparison to the Feedback Vertex Set (FVS) control method

Methods of control based on the structure of the network have been developed by several groups. For example, Zañudo et al. [58] proposed to control a set of nodes intersecting all feedback loops in the network (Feedback Vertex Set) plus the source nodes of the network to attain controllability of the network (other groups have suggested using the feedback vertex set as a control target [30,12]). We used the code provided in https://github.com/yanggangthu/FVS_python to approximate a minimal Feedback Vertex Set presented in Figure 1. This yielded the set of nodes {TP53, E2F1, EGFR, RB1, GRB2, CyclinE1} which is a much larger set of nodes to control. It should be remarked however, that the control goals presented in [58] are more general than our narrow control goal of destroying the existing proliferative steady states.

The method presented in this manuscript allows for more targeted and specific control strategies since we are taking the dynamics of the network into consideration. For example, we encoded our problem as destroying the existing proliferative steady states in which case, we observed control sets of size one. Our method can also be implemented as avoiding solutions to the system {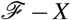, Proliferation − 1} to find controls such that no proliferative state exists.

Some methods of control using the dynamics of networks include methods such as *stable motifs* for guiding the network towards desired attractors or away from undesired attractors [54,55], using the concepts of Boolean canalization, [5,31], and using the concept of the logical domain of influence [52]. To the best of our knowledge, none of these methods have been implemented for general multistate systems. Interestingly, the concept of stable motifs has recently been generalized for multistate networks [13].

### 3.5 Recovering the iron export capability of breast epithelial cells

Iron is an essential metabolite for eukaryotic cells. Iron is necessary for heme biosynthesis, iron-sulfur cluster generation, and acts as a co-factor in several cellular processes such as DNA replication. It is well accepted that iron metabolism is deregulated in several cancers [46,29]. In particular, low expression of ferroportin, the only known non-heme associated iron exporter in mammalian cells, is associated with poor prognosis in breast cancer [34]. In [3], the authors present a ternary (each variable has three states) mathematical model of how iron metabolism interacts with oncogenic pathways (See Figure 2 and Table 3).

**Fig. 2:**
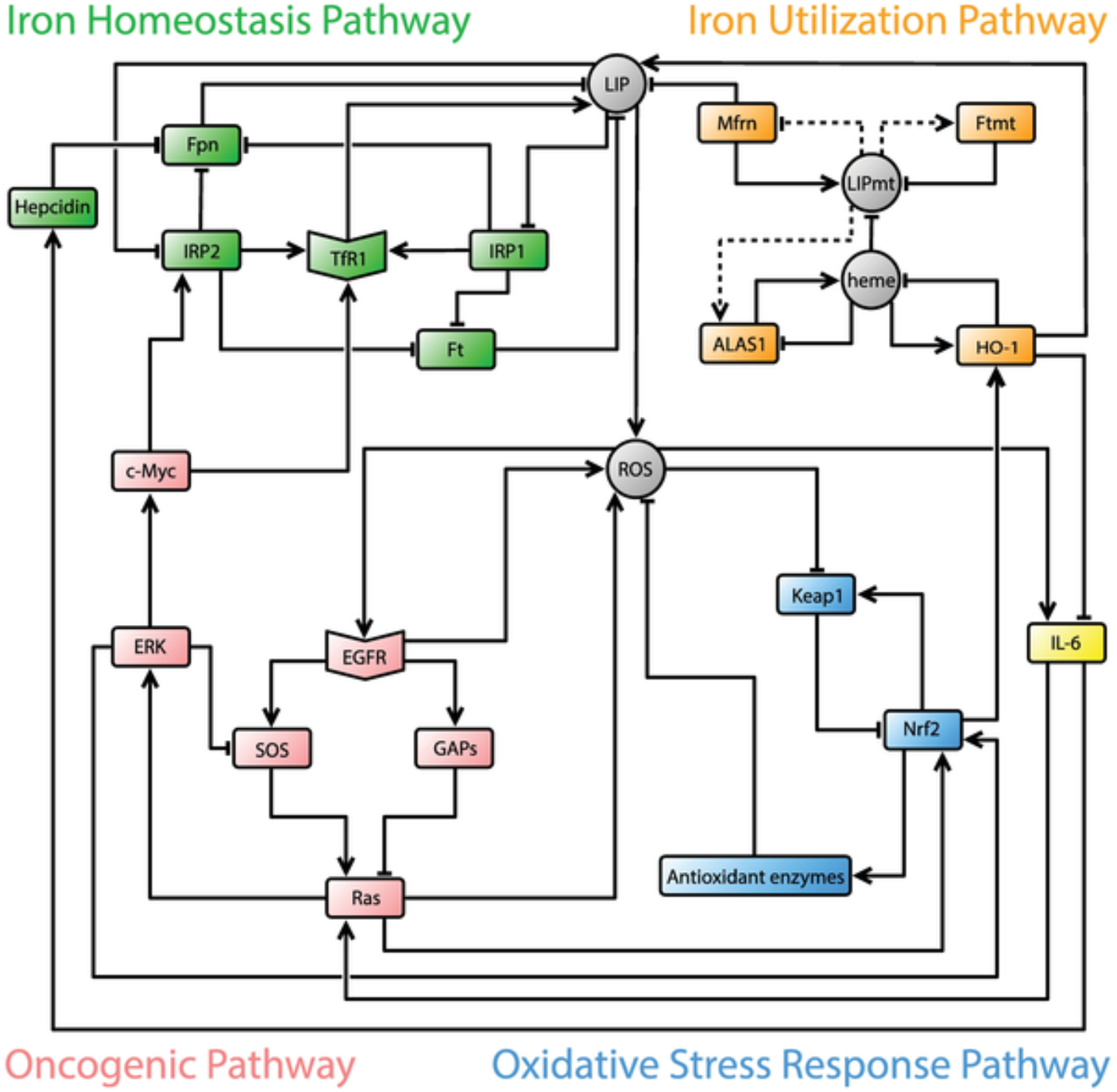
Intracellular iron network published in [3].

**Table 3:**
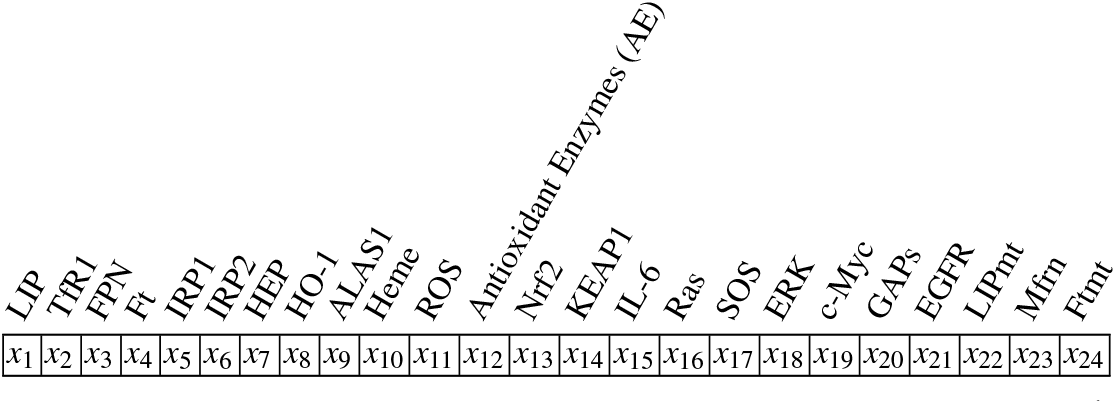
Corresponding variables for the nodes in Figure 2

The model presented in [3] predicts that over-activation of RAS leads to low expression of the iron exporter ferroportin. As previously remarked, low expression of ferroportin correlates with poor prognosis. Hence, regaining basal ferroportin levels might prove beneficial to reduce cell proliferation and, thereby, tumor growth, as observed in mice [34].

We thus encode our control problem as avoiding steady states where ferroportin expression is low. To do this, we allow downregulation of the nodes that are not part of the oncogenic pathway (RAS, SOS, ERK, c-MYC, GAPs, EGFR). In other words, we allow downregulation of the following nodes.

NODE= {LIP, TFR1, FT, IRP1, IRP2, Hepcidin, H0-1, ALAS1, Heme, ROS, AE, Nrf2, Keap1, IL6, LIPmt, Mfrn, Ftmt}.

We encode the control problem as avoiding solutions to the system encoded by the ideal (*F − X*, *x*_3_) where *x*_16_ = 2 (RAS is overexpressed, see Table 3). We note that 10 computation of the generators of the ideal *I* = (*F* − *X*, *x*_3_) took an average of 10.5875s with a standard deviation of 0.2904297, including the overhead of starting the Docker Macaulay2 image.

One of the generators is *u*_6_ * *u*_15_, that is, simultaneous knock-down of IRP2 and interleukin-6. After setting *u*_6_, *u*_15_ to one, that is knocking down IRP2 and interleukin-6, and recomputing the steady states, we get exactly one steady state with ferroportin =1, as desired. That is, our model predicts that knocking down IRP2 and interleukin-6 induction of hepcidin will restore the ability of the cell to export iron. This leads to a hypothesis of whether recovering the ferroportin expression of cancerous cells by knocking down interleukin-6 and IRP2 could lead to cell cycle arrest, similar to iron-chelators [53]. In fact, ferroportin overexpression in prostate cancer cells has shown some interesting results such as induction of autophagy and p21 overexpression [9].

It should be remarked that the original model of [3] contains a continuity condition, e.g. a state cannot change by more than one unit at each time step. In the appendix, Section 7.3, we show that we can remove this condition if the only thing we are interested in is steady states. In this way, it is easier to interpret controls straight from the interaction network.

## 4 Discussion

Encoding discrete dynamical systems as polynomial dynamical systems (PDSs) offers a rich toolbox for the analysis of such models. For example, the computation of steady states and cycles can be encoded as a computation of all the solutions to a system of polynomial equations over a finite field [48], and does not require simulations. Moreover, any discrete dynamical system as defined in this manuscript can be encoded as a PDS, and thus the PDS framework offers an encompassing and general framework for encoding discrete models of biological systems. We previously showed how tools of computational algebra can be applied to find control strategies in Boolean networks [32].

Although multistate systems can be converted into Boolean networks [10], the conversion adds new artificial nodes, and thus models might lose their intuitiveness or become more computationally expensive to analyze. In this manuscript, we have presented a method for extending mixed-state networks into a multistate system in a natural way that preserves steady states. Namely, we convert the system to a PDS over a finite field. We also present control methods based on computational algebra that can generate new steady states, destroy existing steady states, or avoid regions in the state space. Importantly, our control strategies allows for the targeting of both nodes and edges of the wiring diagram.

We also note that, in theory, computing the Gröbner basis for a system of polynomial equations can be computationally expensive (with doubly exponential complexity). However, for many biological systems, computing the Gröbner basis can be achieved in a reasonable time [18] and the computational cost does not seem to correlate with the size of the network but with the average connectivity [47].

Although the methods presented here focus on steady states and synchronous updating schedules many biological systems are modeled with asynchronous schedules and stochastic methods. Developing efficient algebraic methods for the control of asynchronous and stochastic multiscale models presents a rich opportunity for the development of methods more widely applicable to biological systems. We will explore computational algebra methods for general updating schedules in the future.

## 5 Conclusion

In many settings in biology and, in particular, in biomedicine, the ultimate goal of an investigation is the solution of a control problem, and mathematical modeling can be a helpful tool in this endeavor. Modeling dynamic biological networks using systems of ordinary differential equations has the advantage that the modeler has ready access to well-developed rigorous mathematical control theory tools. These work well, when enough quantitative information is available. Some control problems, however, such as the ones considered here, are of a more qualitative nature, such as modifying a network as to change its steady state structure in specified ways. These are not as well suited to a control approach based on differential equations, but fit naturally into a discrete modeling framework. There are now several rigorous approaches to control for discrete models that collectively allow a rigorous solution of a range of control problems. This paper adds another methodology to this collection, based on the principle of representing networks through a collection of polynomials over a finite field, which makes available the algorithms, software, and mathematical tools of computational algebra and algebraic geometry for the solution of a wide range of related problems. The algorithm in this paper makes this approach available for general (deterministic) discrete dynamic networks.

## 6 Supplementary Materials

All scripts used to generate the data in this manuscript can be found in the first author’s github repository https://github.com/luissv7/multistatepdscontrol. The software used for computations of Gröbner bases was a Macaulay2 Docker Image v: 1.14 [15].

## 7 Appendix

In this appendix we denote finite fields with either 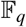 or 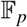, where *p* is assumed to be a prime number while *q* is assumed to be a power of a prime number.

### 7.1 Converting mixed-state models into polynomial dynamical systems

Let *q* be the smallest number which is a power of a prime number such that *q* ≥ |*X_i_*| for all *i*. Consider the finite field 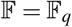.

We can identify 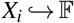 by an injective map *ι_i_* for *i* from 1 to *n*. Let *ι* = (*ι*_1_,…, *ι_n_*). We can now consider the dynamical system **F** as a subsystem of a dynamical system 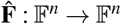 as follows.

Define the map 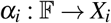 as 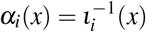 if 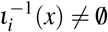 and *α_i_*(*x*) = *c_i_* where *c_i_* ∈ *X_i_* otherwise. Let *α* = (*α*_1_,…, *α*). Now, consider the map 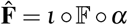.

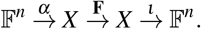

Notice that *α_i_* essentially “crushes” the points in 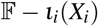 into a constant in *ι_i_*(*X_i_*).

*Example 1* We use a slight abuse of notation for convenience: When we write an integer *m* here, we mean the representative in the particular finite field. Consider the sets *X*_1_ = {0, 1, 2,…, 5}, *X*_2_ = {0,…, 4}.

We embed 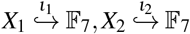 by inclusion. Define *α*_1_, *α*_2_ as follows.

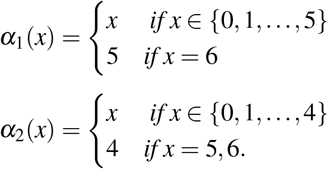

Here *ι* = (*ι*_1_, *ι*_2_), *α* = (*α*_1_, *α*_2_), and 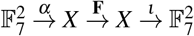.

Veliz-Cuba et al. [48] previously used a similar transformation for a finite field of prime order, 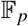, where the elements outside of 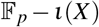 were sent into the “largest” element (*p* − 1). However, in a general finite field 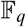, there is no adequate concept of the “largest element”. Notice that 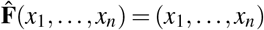 if and only if (*x*_1_,…, *x_n_*) is in the image of *ι* and 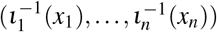 is a fixed point of **F**. In particular, we can now “extend” the discrete dynamical system **F** to a discrete dynamical system 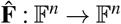 without changing the dynamics of the original system.

### 7.2 An approach for deriving a polynomial dynamical system from a mixed-state dynamical system

A common approach to representing mixed-state dynamical systems is to give Boolean expressions for when a certain node will attain a given value based on the state of the other nodes [56,37]. For example, in the signaling network model presented in [37], the rule for representing how E2F3 attains values 1 or 2 are shown in Table 4.

**Table 4:**
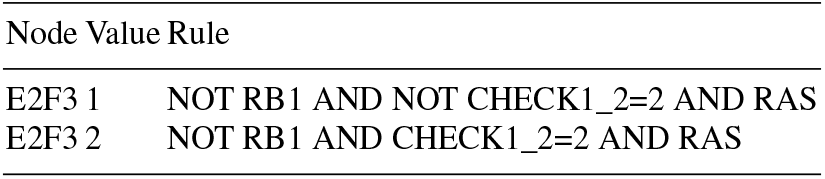
Original equations for E2F3 from [37]

| Node | Value | Rule |
| --- | --- | --- |
| E2F3 | 1 | NOT RB1 AND NOT CHECK1_2=2 AND RAS |
| E2F3 | 2 | NOT RB1 AND CHECK1_2=2 AND RAS |

In the case that some of the variables are Boolean (can only take one of two values), and the other variables are in a set of the same prime cardinality *q*, we can convert to a polynomial dynamical system over 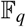. If a variable *x_i_* was Boolean to start with, we replace *x_i_* with 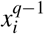. For a variable, *x_i_* that was not Boolean, we can write the polynomial representation by taking advantage of indicators functions 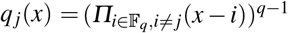 for 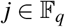. For example, if a variable appears in a Boolean expression as *x_i_* = *j*, then we substitute that variable with 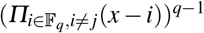. Recall that the operator AND is equivalent to the product over 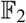, the operator OR is equivalent to the operator (*x, y*) → *x* + *y* − (*x* + *y*) and NOT is equivalent to *x* → 1 + *x*. Over 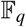, we define *x* AND *y* to be (*x, y*) → (*x* · *y*)^*q*−1^, NOT *x* to be *x* → 1 − *x*^*q*−1^ and *x* OR *y* to be (*x, y*) → −(*x · y*)^*q*−1^ + *x*^*q*−1^ + *y*^*q*−1^.

*Example 2* Consider the update rule for the transcription factor E2F3 from [37], which takes values in the set 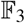, and whose value depends on the nodes RB1, CHECK1_2, and RAS (Table 4). Here, the variables RB1 and RAS were Boolean variables, so we first substitute them with RB1^2^ and RAS^2^. We then apply indicator functions for variables that were not Boolean. For example, CHECK1_2=2 now becomes *q*_2_ (CHECK1_2) where *q*_2_(*x*) = *x* + 2 · *x*^2^.

The final polynomial equation can now be formed by adding the individual functions together, times their respective value.

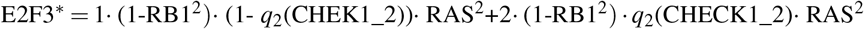

**Table 5:**
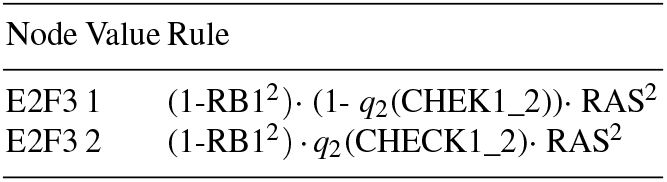
The results of applying the conversion rules to the rules in Table 4

### 7.3 Continuity condition and steady states

The continuity condition is a restriction that the state of each variable does not change by more than one unit at each time step (see *e.g*. [3] for details). Intuitively, the continuity condition represents that a biological quantity cannot suddenly go from high to low (or low to high) without reaching an intermediate step. Here we show that the continuity condition on polynomial dynamical systems used in [3] does not change steady states.

Fix a prime *p* and consider the finite field 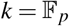. Fix the notation

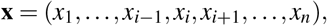

and let **F**_*i*_ := *f_i_*(**x**). We will always assume that the representative for *x_i_* is in the set {0, 1, ⋯, *p* − 1}.

We will say that *f*: *k*[*x*_1_, ⋯, *x_n_*] → *k^n^* is continuous if 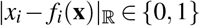 for 0 ≤ *x* ≤ *p* − 1, 1 ≤ *i* ≤ *n*.

Let

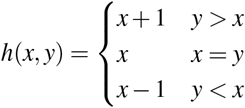

Any PDS **F** : *k^n^* → *k^n^* can be made continuous by considering 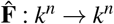 where 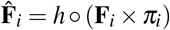 where *π_i_* is the projection onto the *ith* coordinate.

**Theorem 1** *Let* **F** : *k^n^* → *k^n^ be a polynomial dynamical system over a finite field k and let* 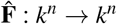 *be the polynomial dynamical system where the continuity condition has been applied to* **F**. *Then the set of fixed points of* **F**, *FIX*(**F**) *is equal to* 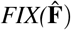

*Proof* Let *x* ∈ FIX(**F**), *π_i_*: *k^n^* → *k* be the projection onto the *i − th* coordinate.

Notice 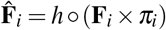. Then 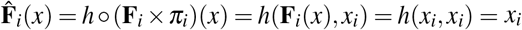.

Now, if 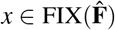, we have *h*(**F**_*i*_(*x*),*x_i_*) = *x_i_* for all *i*. This can only happen if *x_i_* = **F**_*i*_(*x*) for all *i*.

As a result, we have 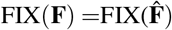.

## Acknowledgements

Sordo Vieira is partially supported by The National Institutes of Health grant no. 1R01AI135128-01. Laubenbacher is partially supported by The National Institutes of Health grants no. 1R01AI135128-01, 1U01EB024501-01, and 1R01GM127909-01.

## Notes

#### Summary of Updates

1. We updated section 2.2.1, where we simplified the encoding of the controls. 2. We added two new sections, Sections 3.3 and 3.4. In these sections we compared our method to other existing methods. 3. We improved the provided code and its efficiency.

https://github.com/luissv7/multistatepdscontrol

## References

1. Albert, R., Thakar, J.: Boolean modeling: a logic-based dynamic approach for understanding signaling and regulatory networks and for making useful predictions. Wiley Interdisciplinary Reviews: Systems Biology and Medicine 6(5), 353–369 (2014)

2. Chaouiya, C., Remy, E., Thieffry, D.: Qualitative Petri Net Modelling of Genetic Networks, pp. 95–112. Springer Berlin Heidelberg, Berlin, Heidelberg (2006)

3. Chifman, J., Arat, S., Deng, Z., Lemler, E., Pino, J.C., Harris, L.A., Kochen, M.A., Lopez, C.F., Akman, S.A., Torti, F.M., et al.: Activated oncogenic pathway modifies iron network in breast epithelial cells: A dynamic modeling perspective. PLoS computational biology 13(2), e1005352 (2017)

4. Choi, M., Shi, J., Jung, S.H., Chen, X., Cho, K.H.: Attractor landscape analysis reveals feedback loops in the p53 network that control the cellular response to dna damage. Sci. Signal. 5(251), ra83 (2012)

5. Correia, R.B., Gates, A.J., Wang, X., Rocha, L.M.: Cana: A python package for quantifying control and canalization in boolean networks. Frontiers in physiology 9 (2018)

6. Cox, D., Little, J., O’shea, D.: Using algebraic geometry, volume 185 of graduate texts in mathematics (1998)

7. Cox, D., Little, J., OShea, D.: Ideals, varieties, and algorithms: an introduction to computational algebraic geometry and commutative algebra. Springer Science & Business Media (2013)

8. Dang, C.V., Reddy, E.P., Shokat, K.M., Soucek, L.: Drugging the’undruggable’cancer targets. Nature Reviews Cancer 17(8), 502 (2017)

9. Deng, Z., Manz, D.H., Torti, S.V., Torti, F.M.: Effects of ferroportin-mediated iron depletion in cells representative of different histological subtypes of prostate cancer. Antioxidants & redox signaling 30(8), 1043–1061 (2017)

10. Didier, G., Remy, E., Chaouiya, C.: Mapping multivalued onto boolean dynamics. Journal of Theoretical Biology 270(1), 177–184 (2011). DOI https://doi.org/10.1016/j.jtbi.2010.09.017. URL http://www.sciencedirect.com/science/article/pii/S0022519310004911

11. Espinosa-Soto, C., Padilla-Longoria, P., Alvarez-Buylla, E.R.: A gene regulatory network model for cell-fate determination during arabidopsis thaliana flower development that is robust and recovers experimental gene expression profiles. Plant Cell 16(11), 2923–39 (2004). DOI 10.1105/tpc.104.021725

12. Fiedler, B., Mochizuki, A., Kurosawa, G., Saito, D.: Dynamics and control at feedback vertex sets. i: Informative and determining nodes in regulatory networks. Journal of Dynamics and Differential Equations 25(3), 563–604 (2013)

13. Gan, X., Albert, R.: General method to find the attractors of discrete dynamic models of biological systems. Physical Review E 97(4), 042308 (2018)

14. Gates, A.J., Rocha, L.M.: Control of complex networks requires both structure and dynamics. Scientific reports 6, 24456 (2016)

15. Grayson, D.R., Stillman, M.E.: Macaulay2, a software system for research in algebraic geometry. Available at http://www.math.uiuc.edu/Macaulay2/

16. Hanahan, D., Weinberg, R.A.: The hallmarks of cancer. cell 100(1), 57–70 (2000)

17. Hanahan, D., Weinberg, R.A.: Hallmarks of cancer: the next generation. cell 144(5), 646–674 (2011)

18. Hinkelmann, F., Brandon, M., Guang, B., McNeill, R., Blekherman, G., Veliz-Cuba, A., Laubenbacher, R.: Adam: analysis of discrete models of biological systems using computer algebra. BMC Bioinformatics 12, 295 (2011). DOI 10.1186/1471-2105-12-295

19. Hinkelmann, F., Murrugarra, D., Jarrah, A.S., Laubenbacher, R.: A mathematical framework for agent based models of complex biological networks. Bull Math Biol 73(7), 1583–602 (2011). DOI 10.1007/s11538-010-9582-8

20. Huang, S.: Gene expression profiling, genetic networks, and cellular states: an integrating concept for tumorigenesis and drug discovery. J Mol Med (Berl) 77(6), 469–80 (1999)

21. Huang, S., Ernberg, I., Kauffman, S.: Cancer attractors: a systems view of tumors from a gene network dynamics and developmental perspective. Semin Cell Dev Biol 20(7), 869–76 (2009). DOI 10.1016/j.semcdb.2009.07.003

22. Ireland, K., Rosen, M.: A classical introduction to modern number theory, vol. 84. Springer Science & Business Media (2013)

23. Kauffman, S.: Homeostasis and differentiation in random genetic control networks. Nature 224(5215), 177–178 (1969)

24. Kauffman, S.A.: Metabolic stability and epigenesis in randomly constructed genetic nets. J Theor Biol 22(3), 437–67 (1969)

25. Klamt, S., Saez-Rodriguez, J., Lindquist, J.A., Simeoni, L., Gilles, E.D.: A methodology for the structural and functional analysis of signaling and regulatory networks. BMC Bioinformatics 7, 56 (2006). DOI 10.1186/1471-2105-7-56

26. Li, R., Yang, M., Chu, T.: Controllability and observability of boolean networks arising from biology. Chaos 25(2), 023104 (2015). DOI 10.1063/1.4907708

27. Lidl, R., Niederreiter, H.: Introduction to finite fields and their applications. Cambridge university press (1994)

28. Liu, J.C., Granieri, L., Shrestha, M., Wang, D.Y., Vorobieva, I., Rubie, E.A., Jones, R., Ju, Y., Pellec-chia, G., Jiang, Z., et al.: Identification of cdc25 as a common therapeutic target for triple-negative breast cancer. Cell reports 23(1), 112–126 (2018)

29. Manz, D.H., Blanchette, N.L., Paul, B.T., Torti, F.M., Torti, S.V.: Iron and cancer: recent insights. Annals of the New York Academy of Sciences 1368(1), 149–161 (2016)

30. Mochizuki, A., Fiedler, B., Kurosawa, G., Saito, D.: Dynamics and control at feedback vertex sets. ii: A faithful monitor to determine the diversity of molecular activities in regulatory networks. Journal of theoretical biology 335, 130–146 (2013)

31. Murrugarra, D., Dimitrova, E.S.: Molecular network control through boolean canalization. EURASIP J Bioinform Syst Biol 2015(1), 9 (2015). DOI 10.1186/s13637-015-0029-2

32. Murrugarra, D., Veliz-Cuba, A., Aguilar, B., Laubenbacher, R.: Identification of control targets in boolean molecular network models via computational algebra. BMC Syst Biol 10(1), 94 (2016). DOI 10.1186/s12918-016-0332-x

33. Naldi, A., Berenguier, D., Fauré, A., Lopez, F., Thieffry, D., Chaouiya, C.: Logical modelling of regulatory networks with ginsim 2.3. Biosystems 97(2), 134–9 (2009). DOI 10.1016/j.biosystems.2009.04.008

34. Pinnix, Z.K., Miller, L.D., Wang, W., D’Agostino, R., Kute, T., Willingham, M.C., Hatcher, H., Tesfay, L., Sui, G., Di, X., et al.: Ferroportin and iron regulation in breast cancer progression and prognosis. Science translational medicine 2(43), 43ra56–43ra56 (2010)

35. Poret, A., Boissel, J.P.: An in silico target identification using boolean network attractors: Avoiding pathological phenotypes. Comptes rendus biologies 337(12), 661—678 (2014). DOI 10.1016/j.crvi.2014.10.002. URL http://dx.doi.org/10.1016/j.crvi.2014.10.002

36. Qiu, Y., Tamura, T., Ching, W.K., Akutsu, T.: On control of singleton attractors in multiple boolean networks: integer programming-based method. BMC Syst Biol 8 Suppl 1, S7 (2014). DOI 10.1186/1752-0509-8-S1-S7

37. Remy, E., Rebouissou, S., Chaouiya, C., Zinovyev, A., Radvanyi, F., Calzone, L.: A modelling approach to explain mutually exclusive and co-occurring genetic alterations in bladder tumorigenesis. Cancer research pp. canres–0602 (2015)

38. Richelle, A., Lewis, N.E.: Improvements in protein production in mammalian cells from targeted metabolic engineering. Current opinion in systems biology 6, 1–6 (2017)

39. Saadatpour, A., Wang, R.S., Liao, A., Liu, X., Loughran, T.P., Albert, I., Albert, R.: Dynamical and structural analysis of a t cell survival network identifies novel candidate therapeutic targets for large granular lymphocyte leukemia. PLoS Comput Biol 7(11), e1002267 (2011). DOI 10.1371/journal.pcbi.1002267

40. Schaaf, G., Hamdi, M., Zwijnenburg, D., Lakeman, A., Geerts, D., Versteeg, R., Kool, M.: Silencing of spry1 triggers complete regression of rhabdomyosarcoma tumors carrying a mutated ras gene. Cancer research 70(2), 762–771 (2010)

41. Shmulevich, I., Dougherty, E.R.: Probabilistic Boolean Networks - The Modeling and Control of Gene Regulatory Networks. SIAM (2010). URL http://www.ec-securehost.com/SIAM/OT118.html

42. Slack, J.M.W.: Conrad hal waddington: the last renaissance biologist? Nat Rev Genet 3(11), 889–95 (2002). DOI 10.1038/nrg933

43. Spencer-Smith, R., O’Bryan, J.P.: Direct inhibition of ras: Quest for the holy grail? In: Seminars in cancer biology. Elsevier (2017)

44. Tan, Z., Zhang, S.: Past, present, and future of targeting ras for cancer therapies. Mini reviews in medicinal chemistry 16(5), 345–357 (2016)

45. Thieffry, D., Thomas, R.: Dynamical behaviour of biological regulatory networks–ii. immunity control in bacteriophage lambda. Bull Math Biol 57(2), 277–97 (1995). DOI 10.1016/0092-8240(94)00037-D

46. Torti, S.V., Torti, F.M.: Iron and cancer: more ore to be mined. Nature reviews Cancer 13(5), 342 (2013)

47. Veliz-Cuba, A., Aguilar, B., Hinkelmann, F., Laubenbacher, R.: Steady state analysis of boolean molecular network models via model reduction and computational algebra. BMC Bioinformatics 15, 221 (2014). DOI 10.1186/1471-2105-15-221

48. Veliz-Cuba, A., Jarrah, A.S., Laubenbacher, R.: Polynomial algebra of discrete models in systems biology. Bioinformatics 26(13), 1637–1643 (2010)

49. Vera-Licona, P., Bonnet, E., Barillot, E., Zinovyev, A.: Ocsana: optimal combinations of interventions from network analysis. Bioinformatics 29(12), 1571–3 (2013). DOI 10.1093/bioinformatics/btt195

50. Waddington, C.H.: The strategy of the genes: a discussion of some aspects of theoretical biology. Allen & Unwin, London (1957)

51. Yadav, V., Zhang, X., Liu, J., Estrem, S., Li, S., Gong, X.Q., Buchanan, S., Henry, J.R., Starling, J.J., Peng, S.B.: Reactivation of mitogen-activated protein kinase (mapk) pathway by fgf receptor 3 (fgfr3)/ras mediates resistance to vemurafenib in human b-raf v600e mutant melanoma. Journal of Biological chemistry 287(33), 28087–28098 (2012)

52. Yang, G., Gómez Tejeda Zañudo, J., Albert, R.: Target control in logical models using the domain of influence of nodes. Frontiers in physiology 9, 454 (2018)

53. Yu, Y., Kovacevic, Z., Richardson, D.R.: Tuning cell cycle regulation with an iron key. Cell cycle 6(16), 1982–1994 (2007)

54. Zañudo, J.G., Albert, R.: An effective network reduction approach to find the dynamical repertoire of discrete dynamic networks. Chaos: An Interdisciplinary Journal of Nonlinear Science 23(2), 025111 (2013)

55. Zañudo, J.G.T., Albert, R.: Cell fate reprogramming by control of intracellular network dynamics. PLoS Comput Biol 11(4), e1004193 (2015). DOI 10.1371/journal.pcbi.1004193

56. Zañudo, J.G.T., Scaltriti, M., Albert, R.: A network modeling approach to elucidate drug resistance mechanisms and predict combinatorial drug treatments in breast cancer. Cancer convergence 1(1), 5 (2017)

57. Zañudo, J.G.T., Yang, G., Albert, R.: Structure-based control of complex networks with nonlinear dynamics. Proc Natl Acad Sci U S A 114(28), 7234–7239 (2017). DOI 10.1073/pnas.1617387114

58. Zañudo, J.G.T., Yang, G., Albert, R.: Structure-based control of complex networks with nonlinear dynamics. Proceedings of the National Academy of Sciences 114(28), 7234–7239 (2017)

59. Zhan, T., Rindtorff, N., Boutros, M.: Wnt signaling in cancer. Oncogene 36(11), 1461 (2017)

60. Zhang, R., Shah, M.V., Yang, J., Nyland, S.B., Liu, X., Yun, J.K., Albert, R., Loughran Jr, T.P.: Network model of survival signaling in large granular lymphocyte leukemia. Proc Natl Acad Sci U S A 105(42), 16308–13 (2008). DOI 10.1073/pnas.0806447105

